# Non-destructive viability assessment of cancer cell spheroids using dynamic optical coherence tomography with trypan blue validation

**DOI:** 10.1101/2024.06.14.598971

**Authors:** Ko Hui Tan, Joel Lang Yi Ang, Alexander Si Kai Yong, Stefanie Zi En Lim, Jessica Sze Jia Kng, Kaicheng Liang

## Abstract

3D cell cultures are widely used in biomedical research for the recapitulation of *in vivo* microenvironments. Viability assessment and monitoring of these intricate conformations remain an open problem as standard cell viability protocols based on colorimetry or microscopy are not directly applicable to intact 3D samples. Optical coherence tomography (OCT) has been explored extensively for subsurface structural and quasi-functional analysis of 3D cell cultures and tissue. Recent studies of dynamic OCT as a source of cellular contrast have found qualitative associations with necrosis in cell spheroids, suggesting potential as a viability marker. We present empirical and validated evidence for dynamic OCT as a quantitative indicator of cell viability in 3D cultures. We analysed over 240 MCF-7 cancer cell spheroids with dynamic OCT and corresponding viability measurements using the trypan blue exclusion assay. Significant effects of common reagents Dimethyl sulfoxide (DMSO) and Phosphate-Buffered Saline (PBS) on OCT readouts were noted. We proposed a regression-based OCT brightness normalisation technique that removed reagent-induced OCT intensity biases and helped improve correspondence to the viability assay. These results offer a quantitative biological foundation for further advances of dynamic OCT as a novel non-invasive modality for 3D culture monitoring.

## 1. Introduction

Three-dimensional (3D) cell culture has revolutionised *in vitro* studies as a more realistic model of *in vivo* cellular interactions than traditional cell monolayers [1]. 3D culture has become indispensable across diverse areas of biomedical research spanning tissue engineering [2], stem cell research [3], clinical pharmacology [4, 5], and even niche fields like space biology [6]. The culturing of cancer cells in dense clumps, known as spheroids, is known to modestly recapitulate *in vivo* tumour biology, while arguably yet to approach the true spatiotemporal complexity of living tumour microenvironments [7].

The power of 3D culture as living models hinges on the accessibility of their real-time physiological status, but assays to obtain such information are inevitably destructive, involving toxic reagents and disruptive interventions. An archetypal example is viability, where the health of the cells within the 3D culture is critical to not only the reproducibility of the model but also longitudinal applications such as cell therapy, transplantation or multi-day/week protocols involving cell differentiation or therapeutics. However, accurate viability monitoring of 3D culture is challenging due to the inaccessibility of cells beneath the outermost layer, hindering the direct application of standard viability protocols based on cell membrane exclusion or metabolism. Reagents for fluorescence/colorimetric viability assays are themselves toxic and also do not reach deeper cell layers, while 3D fluorescence microscopy fails to adequately penetrate moderately sized cultures exceeding 100-200*μ*m diameter [8]. Dissociation or lysis may facilitate the assessment of the culture in its entirety albeit irreversibly destroying the 3D organization with the risk of harming cells [9]. Thus, viability is most commonly predicted via batch sampling from a larger population of cultures, which neglects intra-population variability and fails to characterise the actual samples to be studied or used down-stream.

Non-destructive assays or approaches would, in principle, produce useful readouts with negligible or minimal perturbations to living samples, thereby preserving viability and 3D integrity for longitudinal studies. Non-destructive microscopy based on bright-field illumination or optical phase contrast [10–13] can offer a characterisation of near-transparent 3D samples, but generally assess only structure and provide limited functional information. Optical coherence tomography (OCT) is a non-destructive, non-invasive optical technique best known as an established clinical modality in ophthalmology [14, 15] and cardiology [16], but has also been increasingly touted for cell biology applications due to the potential for safe and label-free volumetric imaging of 3D cell culture, tissue samples and even living animal models [17]. Dynamic OCT (D-OCT) is a variant of OCT that extracts quasi-functional information from samples via signal fluctuations over time and has been shown to differentiate tissue types [18–20] and identify physiological and biological processes [21–23]. The motion contrast detected by the D-OCT technique has been correlated to viability [24–27], but rigorous quantitative validation against standard cell viability assays have been limited [28, 29], hindering widespread adoption by mainstream cell biologists.

We present initial evidence for OCT intensity variation as a quantitative indicator of cell viability in 3D cultures with orthogonal validation through a viability assay. Our findings reveal substantial impacts of Dimethyl sulfoxide (DMSO) and Phosphate-Buffered Saline (PBS) on OCT-based measurements when used as interventional treatments on samples, motivating the development of an adjusted variance metric. Regression analysis demonstrated a strong correlation between OCT intensity variances and cell viability after adjusting for biases induced by these commonly used reagents. This evidence helps bridge the gap between the label-free, quasi-functional capabilities of OCT and the ability for cell biologists to study longitudinal phenomena in a validated experimental pipeline, paving the way for the informed deployment of novel cell monitoring technologies.

## 2. Methods

Spheroid culture is particularly well suited to high-throughput protocols, as a wide range of conditions and interventions may be conveniently tested *in vitro*. The viability of each spheroid or treatment group is an important readout for quantitative comparison. Cell viability may be influenced by various intrinsic and extrinsic factors, including changes in nutrients, oxygen, or stress levels [30]. In 3D cell cultures, a common hypothesis is that cells located near the spheroid core may primarily have decreased access to nutrients and oxygen, suggesting the importance of spatial and subsurface assessment [31]. We aimed to develop a high-content platform [Fig. 1(a)] capable of viability assessment in a non-destructive and non-toxic manner, hence suited for longitudinal studies.

**Fig. 1.**
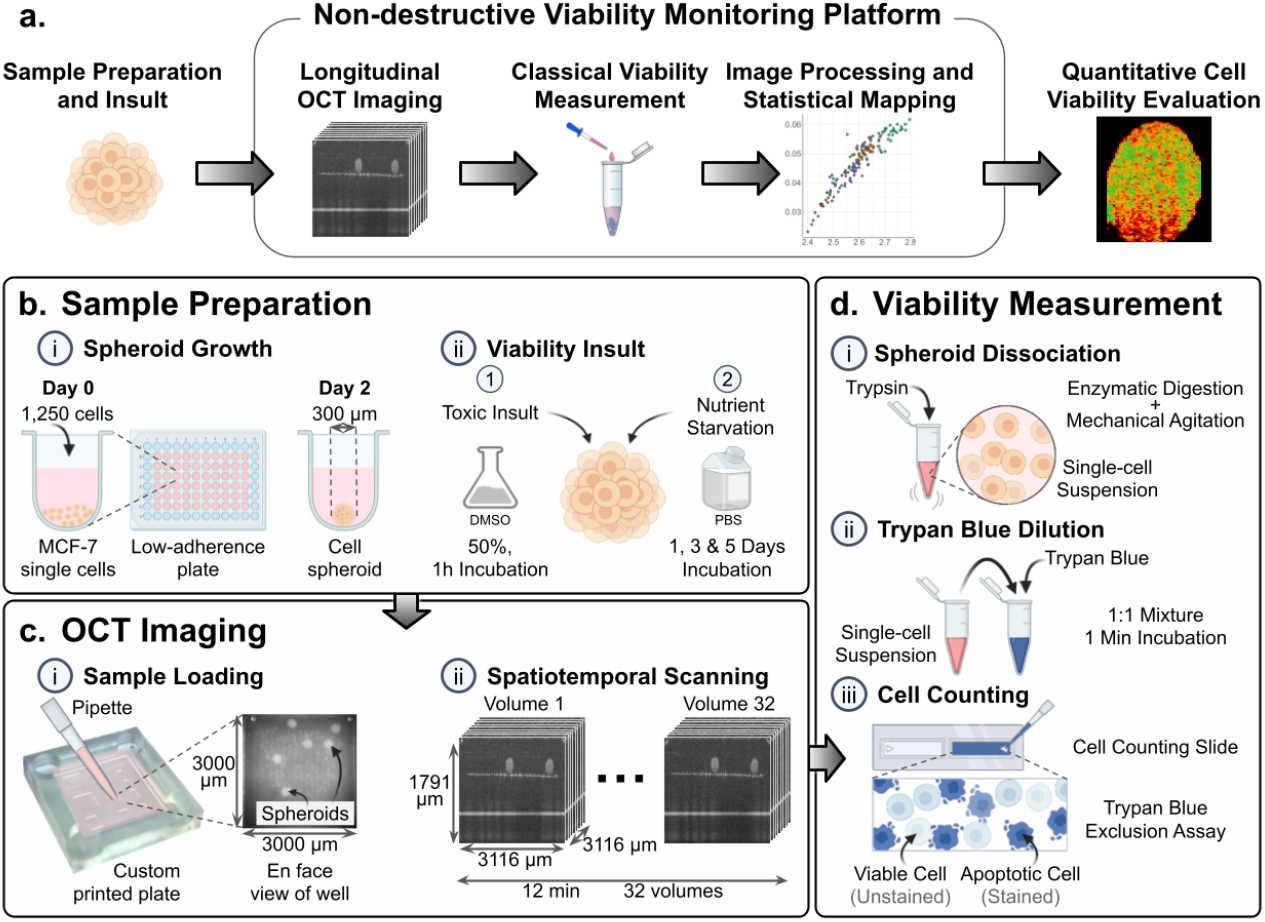
Viability assessment pipeline with biological assay validation. **a**. Proposed workflow for obtaining D-OCT viability mapping for non-destructive imaging. **b**. MCF-7 human breast adenocarcinoma cells were seeded in low-adherence U-bottom plate. Spheroids formed were subjected to viability challenges. **c**. Spheroids were loaded onto custom 3D-printed plate for sequential volumetric OCT scans. Wells of 6 spheroids were imaged over 12 mins to acquire volumes at up to 32 time points. **d**. Trypan blue exclusion assay on dissociated spheroids paired with automated cell counter produced absolute quantitative viability measurements.

### 2.1. Experimental pipeline with biological assay validation

A semi-automated OCT imaging [Fig. 1(a,b,c)] and computational processing pipeline [Fig. 2] streamlined the scanning process for multiple wells containing arbitrary numbers of free-floating spheroids, integrated with a commercially available OCT imaging system (Lumedica OQ LabScope). Numerous spheroids within each well were effectively detected and segmented by K-means clustering, facilitating scalable, high-throughput imaging and analysis. Image registration applied to segmented spheroids from multiple repeat scans acquired at various temporal delays minimised the effects of image drift due to factors such as cell media evaporation, minute environmental perturbations or other fluctuations in the optical signal.

**Fig. 2.**
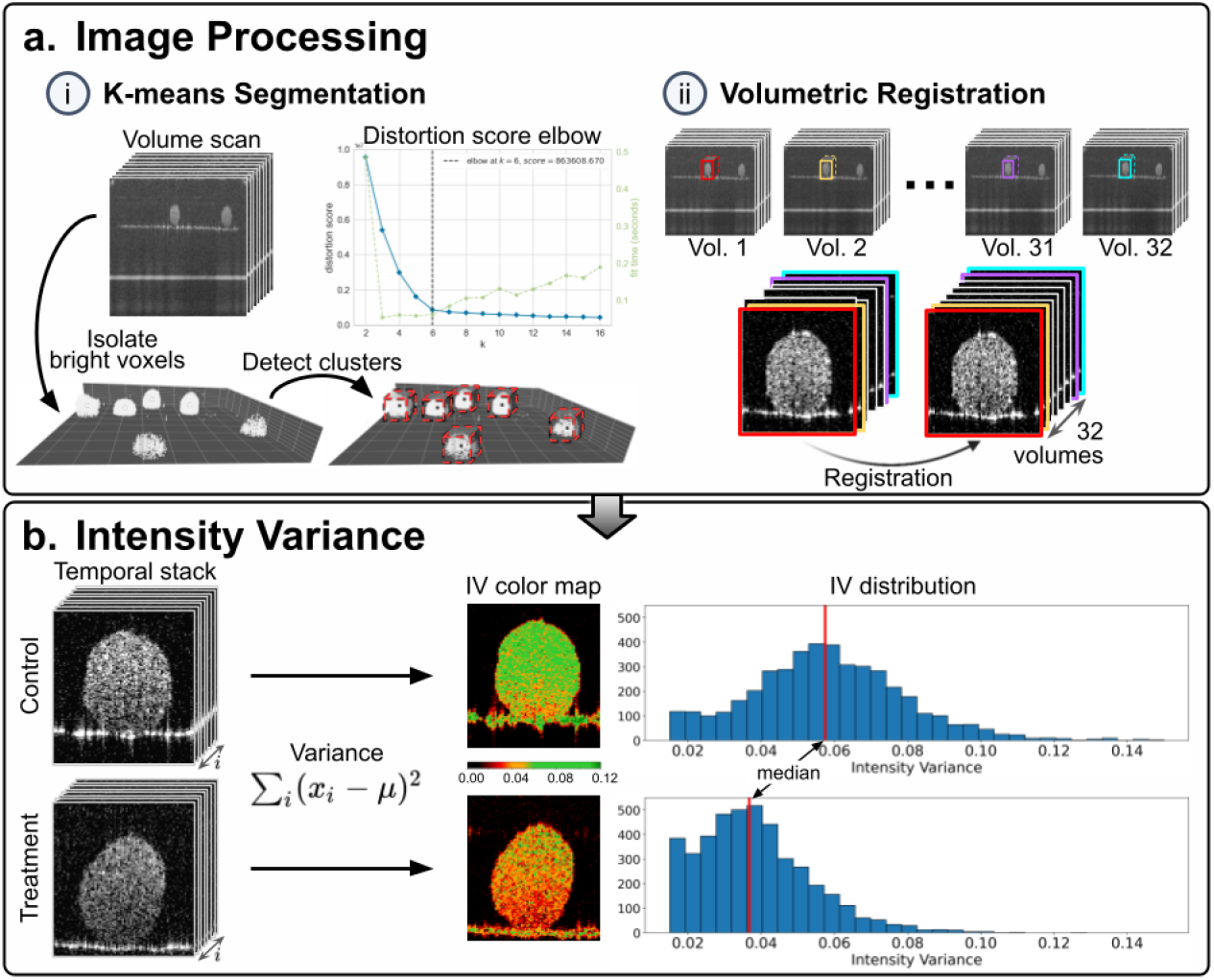
Volumetric image processing for spheroid detection and dynamic motion estimation. **a**. Semi-automated image processing pipeline facilitated rapid analyses on large volumes of spheroids. Instance segmentation and registration improved robustness to micromovements from perturbations during scan. **b**. Intensity variance (IV) served as a quantitative indicator of viability throughout the spheroid. The median IV (MIV) summarises global viability of spheroids for comparison.

Our experimental pipeline included validation of viability [Fig. 1(d)] by a standard laboratory method, where the trypan blue membrane exclusion assay was chosen due to its non-reliance on optical signal for viability readout, thus serving as an orthogonal validation for our optical approach. Immediately following the scan of each sample well, the trypan blue assay was conducted to determine corresponding cell viability [Fig. 1(d)]. The spheroids were gently dissociated into single-cell suspensions using enzymatic digestion with minor mechanical agitation, before dilution with trypan blue. Subsequently, an automated cell counter was employed for high-throughput sampling, ensuring consistent statistical analysis across technical and biological replicates. Due to some loss of cells from the digestion protocol and measurement limits of the cell counter requiring a minimum number of cells, the precision of viability measurements for 300 *μ*m spheroids was restricted to entire wells containing up to six spheroids, such that viability measurement for individual spheroids was not possible in this study.

### 2.2. Intensity-normalised variance metric

OCT signals are sensitive to motion with high spatiotemporal resolution. Temporal (dynamic) fluctuations in OCT signal as a non-invasive readout have been studied under a wide range of name variants intended for different applications from blood flow visualisation to cellular/intra-cellular motion depending on the scan time interval (e.g. speckle variance OCT, dynamic speckle analysis, logarithmic intensity variance, OCT dynamics, etc.). While intensity variation is known to correlate with cellular dynamics, our technology pipeline enabled a close study of dynamic OCT correlations with an orthogonally measured viability readout over different experimental conditions, producing new insights into the biological interpretability of dynamic OCT.

In one possible formulation, the fluctuation signal is defined by the intensity variance (*IV*) over multiple repeated OCT scans (frames or volumes) acquired at a certain interval over time:

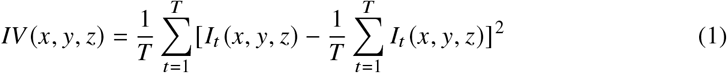

where *I*_*t*_ (*x, y, z*) represents the OCT signal intensity of a voxel at coordinate (*x, y, z*) at time-point *t* for a total of *T* repeated scans.

We used the median of the *IV* [Fig. 2(b)] to estimate the overall variation in an entire spheroid. The median was preferred over the mean due to its inherent robustness to outliers that are common in OCT scans due to reflections.

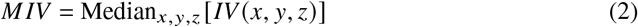

Increasingly, intensity variance is utilised for estimating cell viability, but inconsistent image brightness compromises its accuracy. The dynamic range of a system constrains the observable variance in signal intensity, limiting the detection of subtle fluctuations. In OCT research applications, unanticipated intensity variations can undermine the reliability of D-OCT measurements. These variations may stem from OCT scanning patterns, chemical or mechanical modifications of the sample or its surroundings, and environmental influences. A normalised version of intensity variance has been explored to mitigate this issue [32]. This method, which adjusted deviations from the mean based on mean intensity, was reported to have increased resilience to noise and brightness variations attributable to laser power discrepancies and reflective disturbances. This normalisation reduced the influence of dynamic range alterations on deviation measurements, enhancing measurement consistency across different intensity levels.

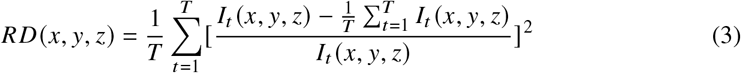

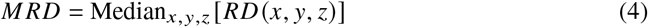

While relative deviation (*RD*) was a convenient formulation that normalised each image voxel with its intensity, preliminary comparisons showed inadequate performance in decoupling variance from overall brightness. We propose an alternative approach for direct normalisation of OCT brightness. The linear relationship between median OCT intensity of spheroids, Median_*x*,*y*,*z*_ [*I*_*t*_ (*x, y, z*)], and *MIV* is first estimated through the slope from a linear fit.

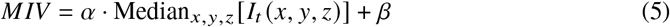

The slope *α* is then used to adjust *IV* to effectively account for the effects of OCT intensity *I* (*x, y, z*). The new quantity, adjusted variance (*AV*), resulting from this normalisation will display a minimal residual linear relationship with OCT intensity.

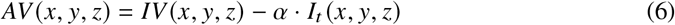

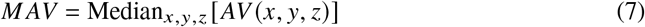

An empirical comparison [Fig. 3] of the three variation forms (*IV*, *RD*, and *AV*) was conducted to establish a good quantitative indicator for cell viability. A strong correlation with cell viability measurements from an established viability assay would suggest the metric’s usefulness as a proxy for cell viability.

**Fig. 3.**
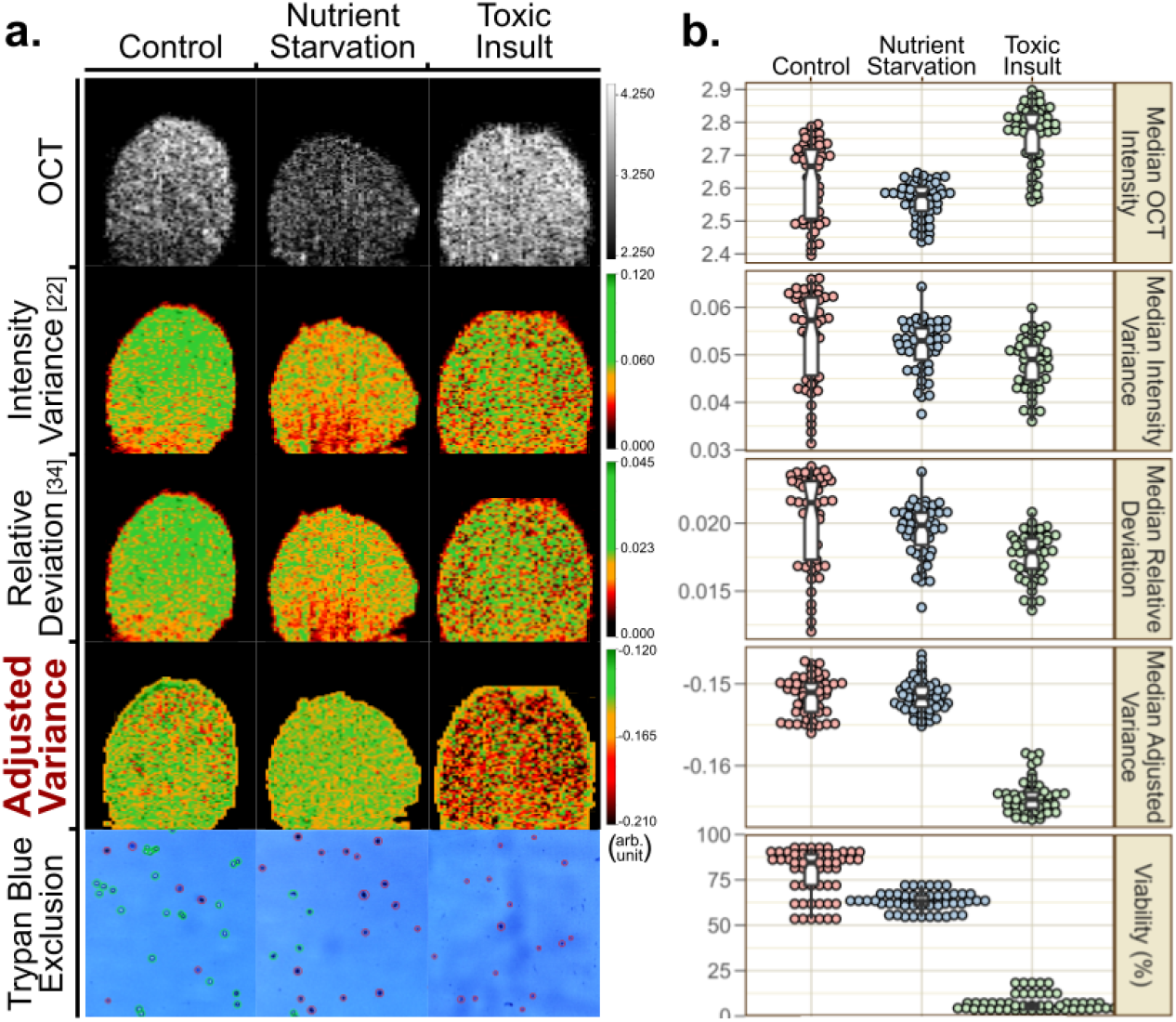
Normalisation techniques to decouple motion from intensity levels. **a**. Sample spheroids demonstrated results of 3 variance formulations on motion estimation. Relative deviation [32] (Eq. 3) exhibits similar patterns to intensity variation [24] (Eq. 1). Our regression-adjusted variance approach removed bias from low intensity near the bottom of spheroids. **b**. Median motion (Eq. 7) distributions of 60 spheroids per viability insult group. MIV exhibits a large range, unable to discern between spheroid groups. Regression adjustment reduces spread and improves separation. Viability distribution obtained via trypan blue exclusion shows high correspondence with MAV distributions.

### 2.3. OCT system and imaging

A commercial spectral-domain OCT system (OQ Labscope 2.0, Lumedica) including a mounted scanner was used for imaging spheroids. It had an A-scan line rate of 13 kHz, a B-scan image rate of 10 Hz, a centre wavelength of 840 nm, and lateral and axial resolutions of 18 *μ*m and 8 *μ*m in tissue respectively as specified by the manufacturer. While these specifications are lower than state of the art OCT prototypes, such a commercial system of low cost and ease of use is most accessible to mainstream users outside of the OCT research subfield and is thus of interest for the present study’s objectives. The horizontal, vertical, and in-plane pixel dimensions were sampled at 256 × 256 × 256 px, resulting in a lateral pixel separation of 12.18 *μ*m and an axial separation was 6.996 *μ*m. Custom OCT imaging plates were 3D printed (Formlabs), with 3 × 3 × 0.5 mm well for spheroid imaging. The width of the imaging well corresponds to the imaging dimensions.

Imaging plates were designed to minimise reflection and output high-quality images by adjusting parameters such as colour of print material, printing resolution and angle, and curing duration. The focal position of the tunable lens in the scanner was adjusted in the system software before each scan to ensure the highest resolution. Spheroids were pipetted onto custom OCT imaging plates filled with media, with 6 spheroids per well. The number of spheroids was optimised to facilitate ease of handling and processing. The same imaging medium (RPMI media) was used to image all spheroid groups. Spheroids were subjected to OCT imaging, with 32 volume scans (C-scan) across 10 min. Custom imaging code enabled automated imaging of spheroids.

### 2.4. Spheroid culture, treatment and validation

Human breast adenocarcinoma epithelial MCF-7 cell line was purchased from the American Type Culture Collection (ATCC). Cells were cultured in RPMI Medium 1640 (22400089, Gibco) supplemented with 10% fetal bovine serum (FBS-HI-12A, Capricorn Scientific), 100 U/mL penicillin and 100 *μ*g/mL streptomycin (15140-122, Gibco) in 5% CO_2_ at 37°C. Formation of spheroids was induced by seeding MCF-7 cells at 750–1250 cells/well in 96-well U-bottomed ultra-low attachment plates (174929, Life Technologies) with shaking. 300 *μ*m spheroids were formed after 2 days. The size of spheroids was estimated with an inverted brightfield microscope (Eclipse Ts2, Nikon).

Samples for the ’live’ spheroid group were obtained from the plate directly without any treatment. Two ’dead’ spheroid groups were prepared with separate interventions known to negatively affect viability - incubation with Dimethyl Sulfoxide (DMSO), or incubation with PBS as a form of nutrient starvation. DMSO has extensive utility in cell biology, including its role as a cryoprotectant for the preservation of cellular specimens [33], as well as a promoter of stem cell differentiation for organoid culture [34]. Although DMSO may induce cell toxicity by apoptosis due to plasma membrane pore formation even at low concentrations [35], it remains widely used in cell biology due to its versatility. Access to nutrients is essential in maintaining cell health. Due to inadequate nutrient diffusion in 3D cultures, particularly in intestinal and brain organoids [36], investigating the effect of nutrient starvation on spheroid viability is biologically relevant. Samples for the ’dead’ spheroid group with toxic intervention was subjected to treatment with 50% DMSO for 1 hour before washing with 1× PBS twice. Samples for the dead spheroid group with nutrient starvation intervention was obtained by substituting media with 1× PBS and left to incubate for 1 day. Each group had 60 spheroids. For the nutrient starvation timepoint experiment, 1× PBS was substituted for 1 day, 3 days and 5 days before imaging, with ∼ 20 spheroids per group. Spheroids were collected from the imaging plates following OCT imaging, and media was replaced by 20 uL 0.25% Trypsin-EDTA (25200056, Gibco). Spheroids were incubated at 37°C for 30 min, for enzymatic digestion into single cells. Mechanical agitation (brief pipetting and vortexing, 1 min each) enabled complete dissociation of spheroids. The cell suspension was mixed 1 : 1 with the Trypan Blue solution (15250061, Gibco) before cell counting (LUNA-II, Logos Biosystem). Viability values were obtained from the machine readout.

### 2.5. K-means object detection

Spheroid detection and locating were performed on the initial volume scan and then applied to subsequent scans in the sequence. Thresholding was employed to isolate bright voxels, capturing spheroids as well as the well’s walls and floor. The walls and floor could be systematically removed due to their consistent vertical and horizontal alignment. The voxels remaining after this filtering process represented the spheroids in the well, which were then identified and located using volumetric K-means clustering. The optimal number of clusters, *K*, corresponding to the number of spheroids, was automatically determined using the elbow method. Although our experiments consistently used 6 spheroids per well, this method is adaptable to varying spheroid quantities within the wells, as long as samples have sufficient spatial separation. We used the scikit-learn [37] implementation of K-means clustering and the Yellowbrick [38] implementation of the elbow method using the Python programming language.

### 2.6. Statistical analysis

Regression analysis, Student’s t-tests, and Pearson correlation coefficient calculation for results demonstrated in Figures 4, 5, and 6 were conducted using R (Version 4.3.2). Levene’s test was conducted before the t-tests to check for equality of variances. Depending on whether the p-value from Levene’s test was statistically significant (*p* < 0.05), either the pooled variance was used to estimate the variance or the Welch (or Satterthwaite) approximation to the degrees of freedom was used. The t-tests were all one-tailed.

**Fig. 4.**
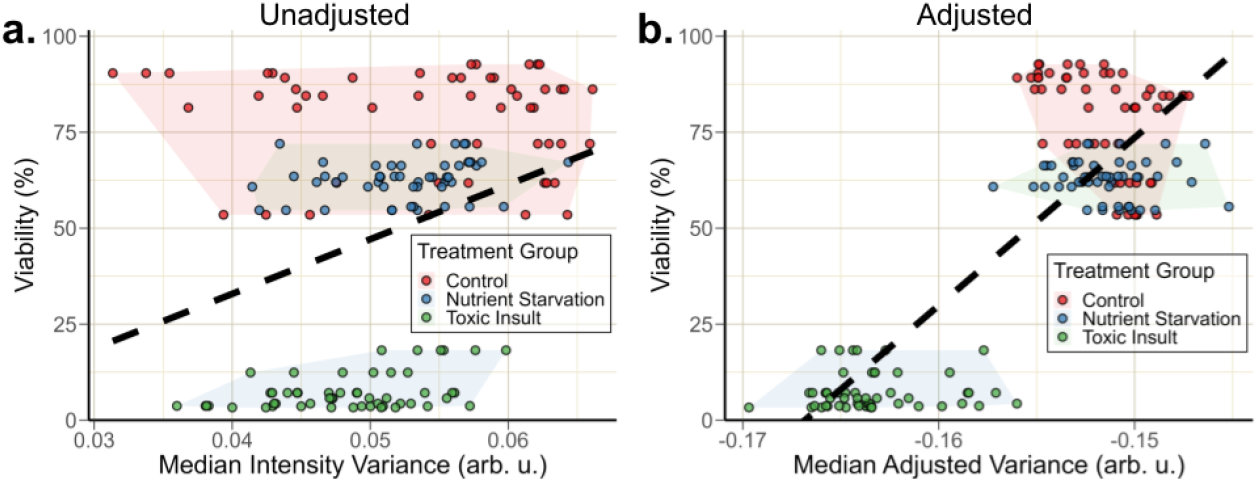
Normalisation of intensity variance (IV) improved linear correspondence to viability measurements. **a**. Analysis showed weak linear relationship between median IV and viability with a Pearson correlation coefficient of 0.305 and an *R*^2^ of 0.093. **b**. After normalization of OCT brightness, median adjusted variance demonstrated stronger linear relationship with a Pearson correlation coefficient of 0.856 and a *R*^2^ of 0.734 and better separability of spheroid groups.

**Fig. 5.**
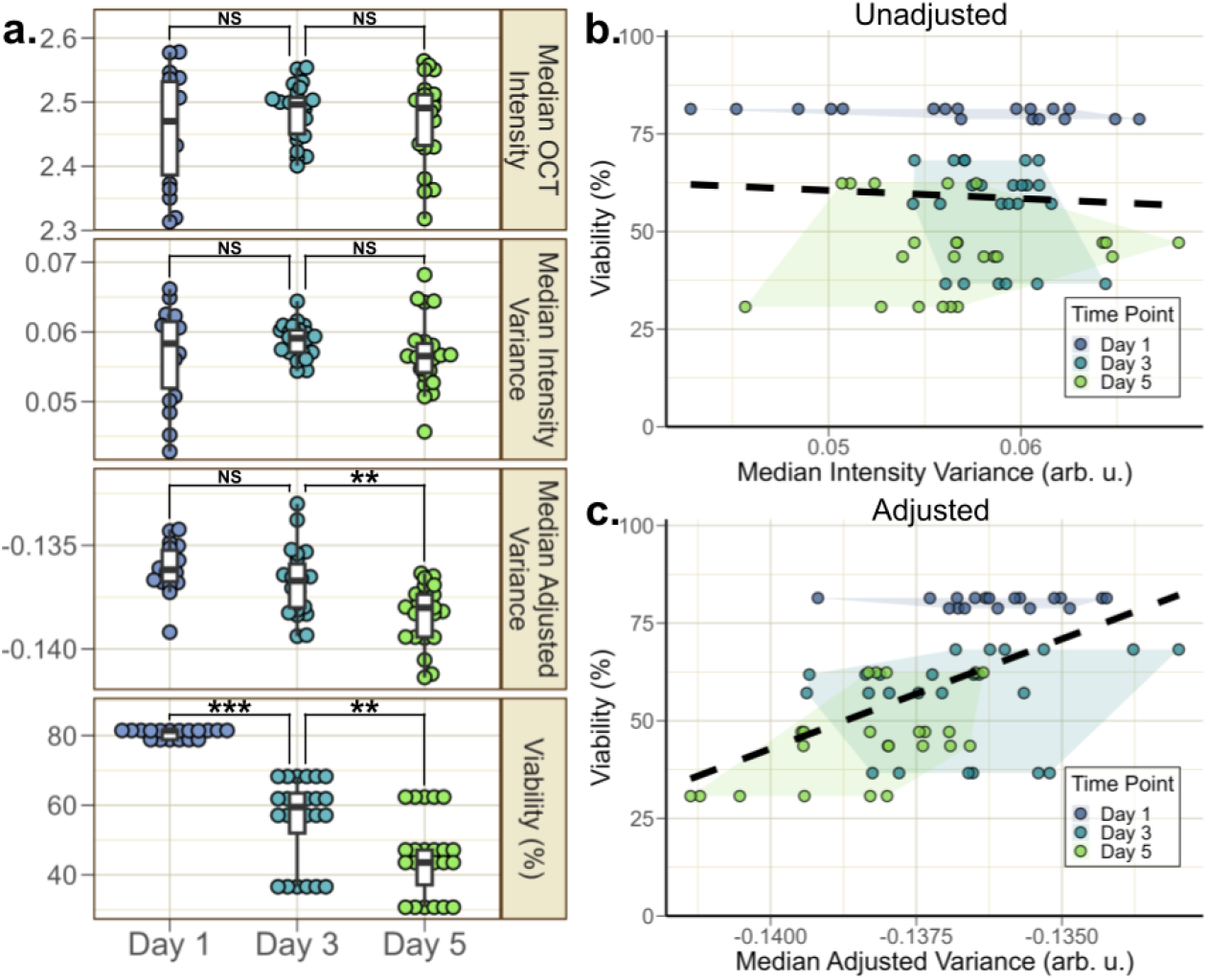
Longitudinal nutrient starvation with normalisation improved separation between groups. **a**. Median intensity variance distributions of ∼ 20 spheroids per starvation time point. Regression adjustment variance by intensity reduced spread and showed distinct groups (*p* < 0.01). Viability distribution obtained via trypan blue exclusion shows close correspondence with MAV distributions (*p* < 0.01). **b**. Analysis showed poor correlation between median intensity variance and viability, with Pearson correlation coefficient of -0.060 and an *R*^2^ of 0.004. **c**. MIV-based investigation required adjustment to distinguish between groups of nutrient-starved spheroids, with Pearson correlation coefficient of 0.548 and an *R*^2^ of 0.300 after adjustment.

## 3. Results

We used our imaging and assessment pipeline to validate practical use cases of non-destructive viability monitoring, while also studying the robustness of optical viability metrics. Two strategies with practical relevance to real-world protocols were employed to induce a lowering of the viability of spheroids as a simulated intervention - introduction of DMSO in the cell media, and nutrient starvation by replacing media with PBS. Crucially, the spheroids regardless of the treatments and viability maintained their overall shape and structure, allowing the consistent use of OCT for our analysis. Overlong incubations with non-ideal reagents occasionally led to spheroid dissociation or disintegration indicating drastic viability deterioration and was thus avoided.

### 3.1. Chemical biases on OCT measurements

We aimed to evaluate the hypothesis that cell viability distinctly influences the *IV* and, thus, could be directly inferred from *IV* measurements. Given that signal intensity constrains dynamic range and thus its variance, we assumed that mean or median OCT signal intensity remains largely unaffected by cell viability. To assess this, three spheroid groups were analysed: a healthy control, one subjected to induced toxicity, and another experiencing nutrient starvation. These varied conditions enabled us to examine *IV* ‘s capability as a marker for cell viability.

Initial observations revealed several key points. OCT imaging consistently showed that structural integrity remained intact across all spheroid groups despite differing treatments [Fig. 3(a)], important for OCT comparisons. Furthermore, the treatments appeared to elicit substantial alterations in OCT signal brightness [Fig. 3(b)], suggesting potential optical effects induced by the chemical agents. Referencing our initial assumption of the independence of OCT intensity from viability, these observations implied that certain chemical reagents could influence the sample’s optical properties with measurable differences in optical imaging characteristics when compared to imaging in conventional cell media.

The treatments’ substantial impact on OCT signal intensity reinforced the importance of using formulations for intensity variance that were normalised to intensity. *MIV* and *M RD*, while having differing absolute values, had indistinguishable distribution patterns across the treatment groups; they both demonstrated large spreads, poor separation between the groups, and poor correspondence to viability readouts. *MAV*, however, demonstrated significantly distinct distributions characterised by reduced intra-group variability and more accurately reflected viability readouts [Fig. 3(b)]. The linear fit and separability of the treatment group clusters substantially improved when using *MAV* as the indicator over *MIV* [Fig. 4, Tab. 1]. Pearson correlation coefficient (***r***) rose from 0.305 to 0.856 and the coefficient of determination (*R*^2^) increased from 0.093 to 0.734.

**Tab. 1.**
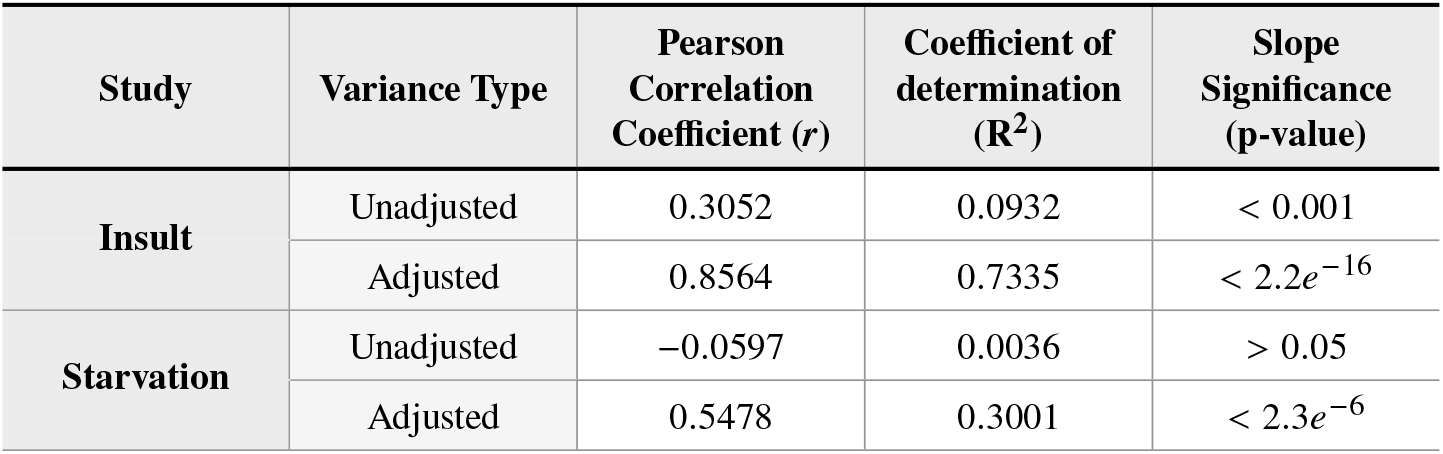
Goodness-of-fit measures of linear models. **I**mprovement in linear fit across all measures using the brightness-normalised variation formulation compared to the standard IV method, indicating enhanced reliability as a proxy for viability. Fit quality varied by experimental setup, showing notably better fit in the multi-insult scenario and poorer fit in the time-point starvation experiment.

Our findings revealed that *IV* exhibited heteroscedasticity; that is, the variability of *IV* across the dataset is not constant but depends on the signal intensity, which in turn depends on the experiment setup and the addition of chemical agents. This heteroscedastic relationship suggests that using a normalised measure of variance, like *AV*, optimised for the specific use case is an important consideration for future reliable adoption of D-OCT for biological inference.

### 3.2. Nutrient starvation longitudinal evaluation

To further our investigation, we conducted a subsequent study focusing on the nutrient-starved spheroid group in an effort to minimise treatment biases. We subjected spheroids to nutrient deprivation and monitored them longitudinally, collecting samples on days 1, 3, and 5. By maintaining the spheroids in PBS throughout the study, we minimised potential OCT brightness biases from chemical interactions. We found no significant change in median OCT intensity across treatment days (*p* > 0.05), substantiating our claim that OCT intensity is influenced by chemical treatment [Fig. 5(a)]. However, there was also no significant change in MIV across treatment days, and the results had a large spread in all groups. Post-processing normalisation with *M AV* effectively improved the separation between spheroid day 3 and day 5 treatment groups, but there was a diminished effect on day 1 and 3 spheroids, likely due to the large MIV range in the day 1 group. Pearson correlation coefficient (***r***) increased from -0.060 to 0.548 and the coefficient of determination (*R*^2^) increased from 0.004 to 0.300 [Fig. 5(b,c), Tab. 1], indicating that the use of *M AV* produced a moderate improvement. With only one treatment group in this investigation, MIV was likely to have limited sensitivity to separate between groups with similar viability and a large spread of data. Hence, further validation of treatment groups may be required to assess their suitability in using optical variance as a tool for viability estimation.

## 4. Discussion

Rapid advancements in the sophistication of 3D models recapitulating physiological and tissue-specific microenvironments are urgently necessitating techniques for non-invasive monitoring of meaningful readouts from these models. The accepted standard of quality control involving the sacrifice of a subset of samples per batch while not interrogating the actual samples to be used is likely unsustainable and inadequate, especially in critical applications such as personalised medicine involving patient-derived material or live transplantation. A non-invasive method of estimating viability could significantly improve the yield and reliability of 3-D culture protocols. OCT technology is already relatively commonplace due to its wide usage from medicine to non-destructive testing, and could be easily repurposed for cell biology applications.

While various metrics including LIV purporting to track OCT dynamics have been suggested, our study uniquely and critically appraises dynamic OCT for its ability to track cell viability in treatment studies explicitly designed to reduce viability with independent validation. The experiments used common interventions namely chemical treatment and nutrient starvation to induce changes in viability, where we found certain reagents to affect OCT image intensity, possibly corroborated by prior work that reported reagents causing alterations in the infrared light spectrum [39–42]. We found that a popular dynamic OCT metric LIV had a substantial correlation to OCT signal intensity, and that an adjusted metric AV accounting for this correlation showed much improved agreement with an orthogonal viability assay. Our experiments simulating viability degradation may not have directly recapitulated the exact mechanisms that occur in natural cell death, and the specific addition of DMSO and PBS to spheroids may have led to changes in sample optical properties that may not be generalizable to other real-world protocols.

The selection of trypan blue exclusion for assessing cell viability was driven by the need for absolute viability measurements using a simple technique that was orthogonal in a sense that it did not rely on optical contrast or involve 3D microscopy. Other possible assays comprised colorimetric or photoluminescence/fluorescence-based assays quantified using a microplate reader device, or 3D fluorescence microscopy using fluorophores that rely on membrane exclusion to indicate live/dead cells. The former group may offer enhanced sensitivity and precision from a quantitative instrument, but only produce absolute measurements and rely on control and background readings from other samples/measurements to produce a percentage readout. The latter group including confocal microscopy offers the ability to perform 3D imaging of membrane exclusion, but has substantially worse imaging depth (<100 *μ*m) than OCT and is thus able to readout from only a very small and superficial fraction of the cells in a spheroid. Trypan blue-based membrane exclusion is a widely recognised and efficient assay producing a percentage readout, compatible with an automated cell counter for systematic interpretation of a bright-field image, and large sample sizes to enhance statistical robustness. The trypan blue assay required spheroid dissociation before viability estimation, involving a combination of enzymatic digestion and mechanical agitation to break up the cell clumps into single cells. We considered this requirement a strength of the approach ensuring a thorough assessment of cells throughout the 3D sample not limited by optical penetration, but such dissociation inevitably led to some loss of cells and might have presumably caused small but measurable changes to cell viability. We considered trypan blue exclusion to be acceptable for our purpose for the rapid tracking of broad viability trends across a sizeable spheroid population and the validation of OCT as a similarly high-throughput approach.

Our study’s OCT system and scanning protocol differed from some prior studies that initially proposed the LIV metric [24, 25]. We utilised a commercially available low-cost spectral-domain OCT system at 800 nm center wavelength (Lumedica), as opposed to lab-developed prototypes that were at 1300 nm and/or used swept laser sources optimizing depth penetration or imaging speed respectively at severe cost to imaging resolution. For applications in cell biology even at high sample volumes, image resolution is almost always vastly preferred by mainstream practitioners over imaging time. There has yet to be a consensus of what may be optimal OCT imaging parameters to derive dynamic OCT measurements of optimised precision, repeatability or other desirable feature. An important imaging parameter and distinguishing feature of our work was the image sampling frequency, in other words the inter-scan time that was used to obtain OCT signal variance. Contrary to previous work computing variance over a set of repeated B-scan frames, in one study as much as 350 B-scan repeats imaged rapidly within 4.48 s [24], we chose to perform repeated volumetric scans, acquiring 32 volumes with a longer inter-scan duration of up to 23 s. Repeat B-scans with intervals in the millisecond range were traditionally preferred in applications deriving image contrast from the dynamics of blood flow, but the time-scale of cellular processes are arguably longer [43]. We chose to use a repeat-volume protocol with significantly longer inter-scan times to avoid biasing towards certain short timescales, which limited sensitivity to particularly fast fluctuations. These differences potentially resulted in significant differences in the sensitivity of OCT measurements.

In this study, we have demonstrated a technical and computational pipeline for quantitative evaluation of cell viability using OCT imaging, compatible with typical interventions used in cell biology, and validation by standard assays. We have found a conditional association between OCT intensity variance and viability, with caveats including a dependence on OCT intensity and a substantial spread in OCT variance that may preclude reliable spot predictions of individual samples. OCT intensity variance alone was an unreliable predictor of spheroid viability, and treatment variables should be considered with the use of this technique. Our method is suitable for comparing sample groups and could be used for quality assurance in comparison to gold standard or positive controls. Further investigations to assess the sensitivity of OCT to changes in viability for different treatments and protocols would aid in assessing OCT as a broadly applicable cell biology tool to monitor 3D cell cultures across a wide range of use cases and scenarios.

## Acknowledgements

This effort was partly conducted at the Institute of Bioengineering & Bioimaging (IBB), Agency for Science, Technology and Research (A*STAR). Figure 1 was adapted from/created with Biorender.com.

## Author contributions

Conceptualisation: KL, JLYA, KHT, ASKY, JSJK

Methodology: JLYA, KHT, KL, ASKY, JSJK

Investigation: KHT, JLYA, SZEL

Visualisation: JLYA, KHT

Supervision: KL

Writing-original draft: KL, JLYA, KHT

## Data and materials availability

Code used for image processing and statistical analysis is available from the corresponding author upon reasonable request. All data needed to evaluate the conclusions in the paper are presented in the paper. Experimental data underlying the results presented in this paper are available from the corresponding author upon reasonable request.

